# A study of the expression of germline-specific paralogs of the ubiquitously expressed NAC complex revealed their association with centrosomes

**DOI:** 10.1101/2022.12.12.520084

**Authors:** Elena A. Mikhaleva, Natalia V. Akulenko, Oxana M. Olenkina, Yuri A. Abramov, Sergey A. Lavrov, Thoomas A. Leinsoo, Galina L. Kogan, Vladimir A. Gvozdev

## Abstract

Protein homeostasis involves the ubiquitously expressed heterodimeric protein NAC (ribosome associated nascent polypeptide complex), consisting of α- and β-subunits. We previously found in *Drosophila melanogaster* the presence of germline-specific paralogous genes for NAC subunits (gNACs): unique for the α-subunit and several highly homologous amplified genes for the β-subunit, indistinguishable by the antibodies used, while they were detected as different phosphorylated isoforms in 2D electrophoresis with immunostaining. The distinct increase in gNAC expression in growing spermatocytes compared with their predecessors, gonioblasts, is accompanied by a clear presence of gNAC in the cytoplasm of dividing meiotic cells and spermatids. In oogenesis, gNAC expression is significantly lower than in the testis, but gNAC accumulation is found at the posterior pole of the mature oocyte, known as the germplasm region responsible for the germ cell formation in the offspring. gNAC-associated ribosomes from mature oocytes are enriched in mRNA sets encoding known germplasm and centrosome proteins. gNAC is also detected in a pericentriolar component of centrosomes: in syncytial dividing nuclei in the early embryos, in primordial germ cells and spermatocytes, including meiotic divisions stage. CRISPR/Cas9-mediated deficiency of the gNAC-alpha subunit causes female infertility. We propose the role of the gNAC phosphoregulatory paralog system as a component of the germline proteostasis network, coordinated with the centrosome functions. Here, we also extended our results by identifying cross-links between ubiquitous and germline-specific NAC-β paralogs, finding an additional case of functional cross-talk between heterodimeric ubNAC and gNAC α-subunit paralogs, which may maintain the robustness of proteostasis and fly fecundity.

## 1 Introduction

The role of global regulation of mRNA translation in germ cell development has been actively discussed (Dodson and Kennedy, 2020; Mercer et al., 2021), but evidences for individual specific factors associated with ribosomes and acting in germline protein homeostasis (proteostasis) remains inconclusive. Here, we present the results of our studies of the developmental expression in *Drosophila melanogaster* of the recently detected germline specific paralogs (gNAC) (Kogan et al., 2022) of the known ubiquitously expressed heterodimeric protein NAC (nascent associated protein complex containing NACα and NACβ subunits). The ubiquitously expressed NAC (hereafter also referred to as ubNAC), binds to ribosomes and interacts with nascent polypeptides, considered an important protein providing cellular proteostasis as a sorting partner for SRP (signal recognition particle) providing specificity for the membrane and secretory protein localization in eukaryotes (Kramer et al., 2019; Hsieh and Shan, 2021; Jomaa et al., 2022).

Germline-specific heterodimeric NAC paralogs (gNACs) have been shown to be encoded by a unique gene for α-subunit and several amplified highly homologous genes for the β-subunit paralogs (Kogan et al., 2022). The germline gNACα and gNACβ paralogs were characterized by the acquisition of the extended intrinsically disordered regions (IDRs) compared with the ubiquitously expressed paralogs (Kogan et al., 2022). Below, for simplicity, we will denote by gNAC the sum of these paralogous proteins including different β-subunits that cannot be differentiated using available antibodies. The expression of gNACs in germ cells will be shown using Ab against β-subunit, which does not distinguish these paralogs in tissue staining, but convincingly demonstrates their germline specificity (Kogan et al., 2017; Kogan et al., 2022).The existence of different degrees of phosphorylation of β-subunit paralogs was shown by 2D electrophoresis analysis (Kogan et al., 2022). Phosphorylation sites preferentially located in the extended intrinsically disordered regions (IDRs) of the proteins known to be prone to posttranslational phosphorylation (Gao and Xu, 2012) were predicted in the amino acid sequences of β-subunits (Kogan et al., 2022).

We performed expanded characterization of gNAC expression profiles in spermatogenesis compared with concise earlier reports (Kogan et al., 2017; Kogan et al., 2022) and new results of their detection in oogenesis and embryos that revealed an unexpected location of gNAC in centrosomes. We found strong upregulation of gNAC in the cytoplasm of primary spermatocytes, which persists in meiosis and spermatid differentiation. Accumulation of gNAC was observed in the posterior pole of the mature oocyte, known as the germ plasm region that is essential in embryos for the formation of primordial germ cells. We traced the redistribution of gNAC granules in the fertilized syncytial embryo to the centrosomes of dividing nuclei and the subsequent presence of gNAC in germ buds and pole cells, primordial germ cells. It was found that both gNAC and the germinal marker, the VASA protein, can be detected in the cytoplasm of primordial germ cells and their precursors migrating to the sites of gonad formation. The importance of the presence of gNAC in embryonic syncytial centrosomes is highlighted by recent results revealing the critical role of centrosomes in the development of pole buds and pole cells and the formation of microtubules involved in the proper distribution of germplasm components in primordial germ cells (Lerit and Gavis, 2011; Fang and Lerit, 2020) and in protein interactions (Nishi et al., 2014; Koike et al., 2020; Rieloff and Skepo, 2020). We speculate that these events may be closely supported by a specific germline proteostasis network involving gNACs as internally disordered proteins carrying phosphosites. The mechanisms and interactions of gNAC isoforms with putative partners remain enigmatic, as well as the role of its putative phosphoregulatory modifications, but the very fact that gNAC was discovered (Kogan et al., 2017), opens the way to unraveling its direct functional interactions in germ cells.

## 2 Results

### 2.1 CRISPR-Cas9 mediated deficiency of a unique gene encoding the α-subunit of gNAC causes female infertility suppressed by transgenic overexpression of the ubiquitously expressed paralog

The presence of the unique gene *CG4415* encoding the α-subunit of gNAC in the *Drosophila* genome allowed its local CRISPR-Cas9-dependent inactivation, which would have been virtually unrealizable for the scattered amplified gene set of paralogous gNAC β-subunits (Kogan et al., 2022). Infertility in flies carrying the canonical single CRISPR-Cas9-mediated *DsRed* donor DNA insertion replacing the *CG4415* gene was not detected (mDsRed line), but female sterility was achieved in the incDsRed line carrying tandem repeat of two full and one partial copy of *DsRed* donor construction (see Methods, Figure S1). The *CG4415* gene overlaps with the intron of the transcript encoded by the gene of unknown molecular function *CG31795* (Flybase), but specific damage the *CG4415* gene function in the incDsRed line has been documented by almost complete suppression of female infertility caused by ectopic expression of the transgene encoding the α-subunit of gNAC under the constitutive actin-5C promoter. We previously found that transgenic overexpression of the gNAC β-subunit can suppress the lethal effect of a deficiency of the ubiquitously expressed β-subunit of ubNAC (Kogan et al., 2022). Cross-talks between these paralogs can be considered as a way to maintain the proteostasis robustness in both somatic and germ cells. At the same time, CRISPR-Cas9-mediated canonic replacement of the *CG4415* gene by a single *DsRed* vector copy preserved fly fertility that can be related to a particular robust resilience of this trait maintained by paralogous or other complicated buffering interactions between different genetic systems. Such explanations have been advanced in the absence of a phenotypic effect caused by the monogenic knockout (Dede et al., 2020).

### 2.2 gNAC expression in oogenesis and embryos

The first reliable presence of gNAC in oogenesis was detected in the cytoplasm of the developing oocyte (stages 4-5), in its periphery or around the oocyte nucleus (Figure 1, A-A’’’). There were no signs of the presence of the gNAC protein in the nurse cells cytoplasm (Figure 1, A’, red asterisk). Mature oocytes are characterized by the accumulation of gNAC and gNACβ mRNA at the posterior pole of the oocyte, at the cortex (Figure 1, B-B’’), in the area of the germplasm necessary for the development of primary germ cells in embryos (Ephrussi et al., 1991; Lasko, 2020). Germplasm is stably attached to the actin cytoskeleton during oogenesis and is released from the cortex after fertilization (Lerit and Gavis, 2011).

**Figure 1.**
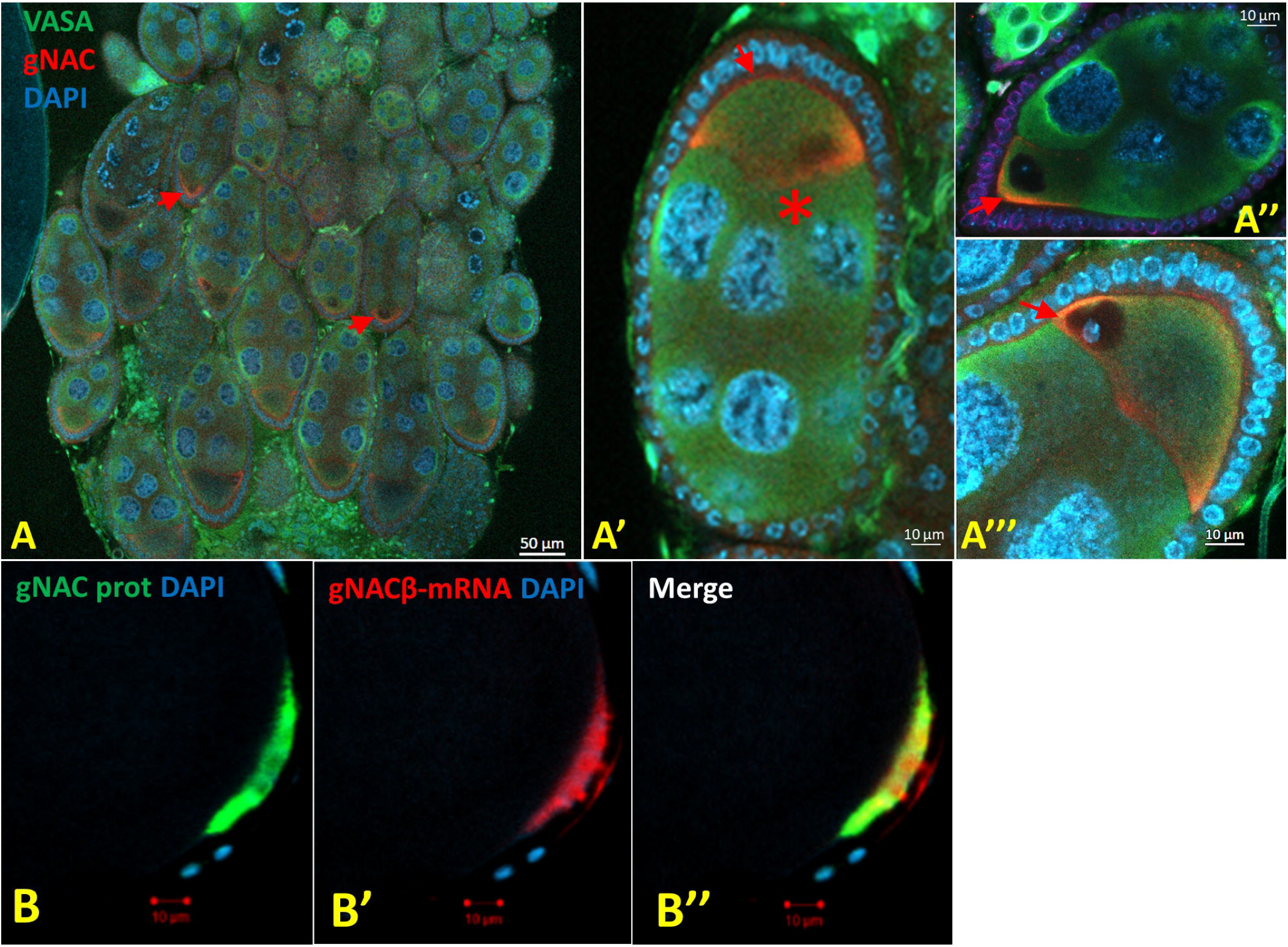
gNAC protein and gNACβ mRNA in oogenesis (A) Overview of ovarioles, onset of gNAC accumulation at the periphery and posterior pole of the developing oocytes (red arrows). (A’) Enlarged egg chamber, stage 4-5 oocyte, gNACβ at the oocyte periphery (red arrow), gNAC absent in the cytoplasm of the nurse cells (red asterisk). (A’’) First signs of accumulation of gNAC at the posterior pole of the oocyte (red arrow). (A’’’) gNAC is located around the oocyte nucleus, stage 4-5 (red arrow). Merged channels images, VASA green, gNAC red, DAPI blue. (B) gNAC protein and gNACβ-mRNA (FISH) at the posterior pole at the cortex of a mature oocyte. gNACβ protein, green (B); gNACβ-mRNA, red (B’); merged image (B’’).

Previously, we found that a major part of cellular gNAC in testis is associated with ribosomes (Kogan et al., 2017). Here, we tested whether there is in ovaries a certain selectivity in the set of mRNAs associated with the gNAC-marked immunoprecipitated ribosomes compared to the individual mRNA distribution in total ribosomes. This proposal was inspired by the widely discussed ideas about the heterogeneity and translational specificity of ribosomes, which can be determined, apart from the specificity of the ribosomal proteins or RNA themselves, by the ribosome associated proteins (Xue and Barna, 2012; Li and Wang, 2020). The RT-PCR analysis of the abundances of individual mRNAs in the sets from the gNAC-associated ribosomes isolated by immunoprecipitation and total ribosomes was performed and revealed up to 10x-20x fold enrichment in mRNAs encoding the known germ plasm proteins (Osk, Vasa, Tud, CycB etc.) and several evolutionally conserved centrosome proteins (Asl, Ana1, mAna2, CP190 etc.) in the gNAC associated ribosomes (Figure 2, see Methods for details) compared to the total ribosomes. At the same time, some mRNAs encoding housekeeping proteins (Adh, αTub, Act) were shown to be randomly distributed between these ribosome samples (Figure 2).

**Figure 2.**
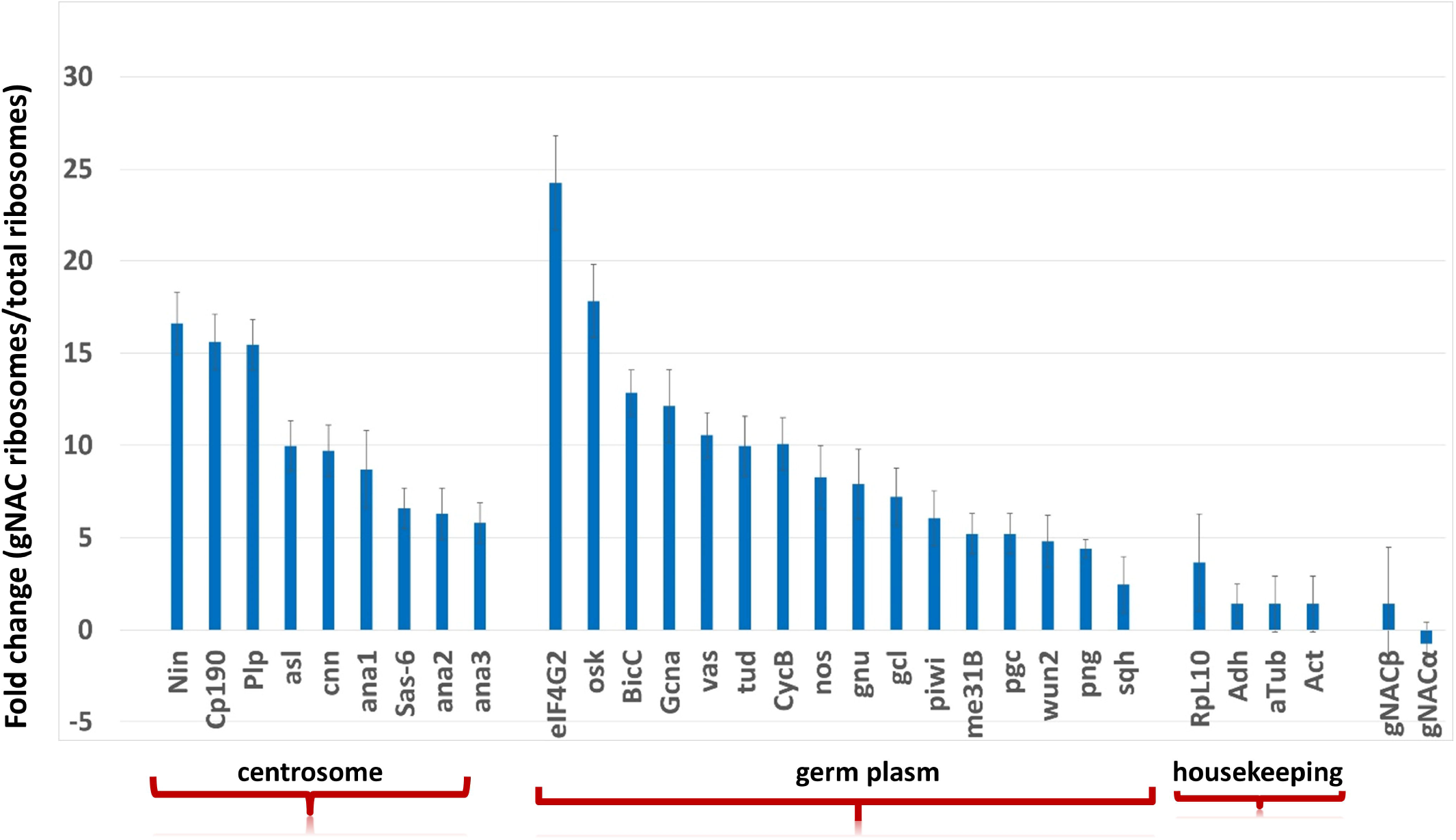
Comparison of mRNA amounts in gNAC-associated ribosomes and total ribosomes in mature oocytes. Contents of mRNAs encoding the germ plasm and centrosome proteins are increased 15 to 25x, respectively, in ribosomes associated with gNAC compared to total ribosomes in ovaries (numeric data are in the Table S1, mRNAs amounts are normalized to *Rpl32* mRNA).

The early stages of embryonic development after fertilization represent the successive high-speed divisions of syncytial nuclei, which at the posterior pole of the embryo form so-called pole buds converted to the pole cells, primordial germ cells (reviewed in (Lehmann, 2016; Lasko, 2020)). To clearly detect centrosomes in syncytial dividing nuclei, we used the stock expressing under endogenous promoter the centriolar Asl-GFP fused protein known to be important to recruit the pericentriolar proteins to centriole forming the pericentriolar material, PCM (Conduit et al., 2014). PCM can be detected using antibody against centrosome protein CP190 known also as a component of insulator complex associated with multiple chromosomal sites as well with the mitotic spindles (Plevock et al., 2015).

In the syncytial cycle 3 (Figure 3, A-A’), the pericentriolar protein CP190 begins to concentrate around Asl-GFP-labeled centrioles, where gNAC accumulation is not yet detectable. Then, in cycles 5-6, some hints of gNAC accumulation (a cloud of gNAC pellets) and PCM labeled with pericentriolar protein CP190 are partially overlap (Figure 3, B-B’). In the cycles 10-12, gNAC granules clearly accumulate around the duplicated centrioles, overlapping with the CP190 protein (Figure 3, C-C’). In cycles 13-14 (Figure 3, F), the blastoderm cells anchored to the embryonic cortex contain cytoplasmic gNAC granules, including apically located centrosomes stained for gNAC (Figure 3, F’’’), which may have a specific cell anchoring function on the cortex (Lv et al., 2021). CP190 spreads from centrosomes to chromosomes to perform the known insulator function (Figure 3, F’). The overlay of gNAC on CP190 flares is clearly visible.

**Figure 3.**
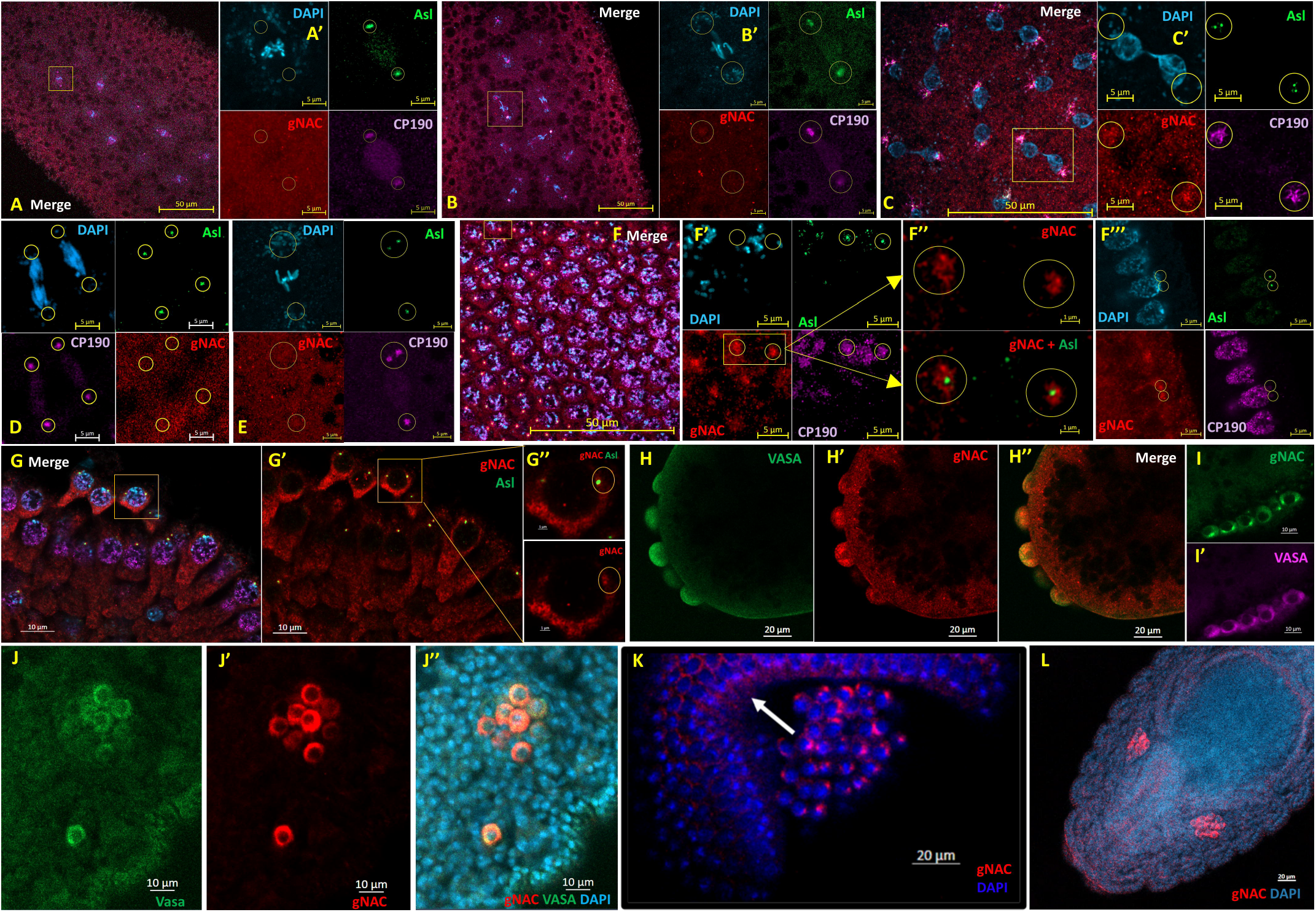
gNAC protein in early embryos (syncytium, pole cells, migrating germ cells). (A) Syncytial division cycle 3, overview of the first nuclear divisions and enlarged image of the division in separate channels, DAPI - blue, Ast-GFP - green; gNAC - red; CP190 - violet (A’). No evidence of gNAC concentration around the centriole, while CP190 is accumulated at centrosomes (encircled). (B) Divisions 5-6, metaphase; onset of gNAC accumulation around centrioles and enlarged image of dividing nuclei in separate channels (B’). (C) Divisions 10-12, merged channels overview and enlarged nuclei (C’). Telophase stage, gNAC staining overlaps with the CP190 flares. (D, E) Divisions 10-12, anaphase (D) and metaphase (E), weak gNAC accumulation in anaphase and metaphase. (F) Blastoderm stage, cycle 13-14, overview of blastoderm nuclei, merged image. (F’), enlarged prophase nuclei, separate channels. Centrosome and nuclear localization of the CP190. Centrosomes contain centriole (green spot) surrounded by PCM with the overlapped regions of gNAC and CP190. (F’’), enlarged part of (F’), yellow rectangle. The distribution of gNAC around Asl-GFP marked centriole is shown. (F’’’), blastoderm nuclei and adjacent duplicated centrioles at the periphery of the embryo; CP190 is localized in the chromatin, gNAC is concentrated around the centrioles. (G) Pole cells, cycle 13, merged channels overview. (G’), separate gNAC (red)+Asl(green) image, (G’’), enlarged pole cell from yellow rectangle area in (G’), accumulation of gNAC around pole cell, centrioles are encircled. (H) Vasa and gNAC in the pole buds. Vasa stained green (H), gNAC is red (H’), (H’’) is a merged image. (I) gNAC accumulation in pole cells centrosomes (I), cytoplasmic location of VASA (I’). (J) gNAC in migrating cluster of primordial germ cells ((J) - Vasa, (J’) - gNAC, (J’’) - merged+DAPI). (K) Primordial germ cell cluster (gNAC staining) during invagination (white arrow). (L) Germline cells are intermingled with somatic cells after lateral migration into compact gonads.

Magnification distinctly shows the position of the centriole surrounded by pericentriolar gNAC material (Figure 3, F’’). A significant cytoplasmic gNAC staining is found in pole buds and pole cells, here cytoplasmic gNAC is also found surrounding the centrosomes (Figure 3, G, H, I). The presence of cytoplasmic gNAC is retained in the cluster of migrating germ cells (Figure 3, J, K) transported during gastrulation (Figure 3, K, white arrow) to the sites of primordial somatic gonadal cells where somatic and germ cells are intermingled forming a pair of gonads (Figure 3, L).

### 2.3 gNAC expression in testis

The incomparably higher expression of NACβ subunit in the testis than in the ovaries is evidenced by the Flybase data and supported by our recent results of the evaluations of the amount of NACβ mRNA encoding gNAC β-subunit (Kogan et al., 2022). A rough evaluation of gNAC presence by immunostaining is in accord with these results. As in the other studies of the spermatocyte specific expressed genes (Lu et al., 2020), we found in primary spermatocytes, in contrast to mitotically dividing precursors, spermatogonia, the upregulated expression of the genes encoding highly homologous gNAC β-subunits, whose expression can be traced to a strong increase in gNAC staining using against its β-subunit Ab, which recognizes several isoforms encoded by highly homologous amplified genes (Kogan et al., 2012) (Figure 4, A, B). A sharp increase of the mRNA presence is also detected in the spermatocytes compared to the spermatogonia at the testis tip (Figure 4, B). A clear level of immunostaining for gNAC is coupled with the presence of germ cell marker Vasa in the cytoplasm of the both spermatogonia and spermatocytes which is conserved in the course of the meiotic divisions and in the cytoplasm of the round and elongated spermatids (see below). No presence of gNAC is detected in somatic supportive cyst cells marked by DAPI stained nuclei (Figure 4, C-C’’). In the primary spermatocytes, no accumulation of gNAC around centrosome is detected (Figure 4, E-E’’). In the dividing meiosis I spermatocytes, the distribution of gNAC granules along the microtubule asters is seen (Figure 4, F-F’’, G-G’’). gNAC staining is distinctly seen in the cytoplasm of round spermatids with single centrioles (Figure 4, H, H’). Cytoplasmic staining’s for gNAC and Vasa are detected in separate compartments of round spermatids (Figure 4, I, I’). In meiosis II, gNAC staining is detected near duplicated centrioles labeled with Asl-GFP (Fig. 4, J) and as the circumference of centriolar microtubules associated with Asl-GFP (Figure 4, K-K‵‵‵). In the beginning of spermatid elongation, the gNAC spot is accumulated at the cytoplasmic distal end of leaf stage spermatids that is conserved during further elongation, at the needle stage. At the same time, gNAC is surrounded the spermatids nuclei, possibly associated along microtubules, known to shape the nucleus during elongation as well at the centriole region nucleated axoneme microtubules (Figure 4 L, L’). Figure 4, M-M’’ demonstrates gNAC distribution (red) along the axoneme microtubules (violet), indicated by yellow arrowheads.

**Figure 4.**
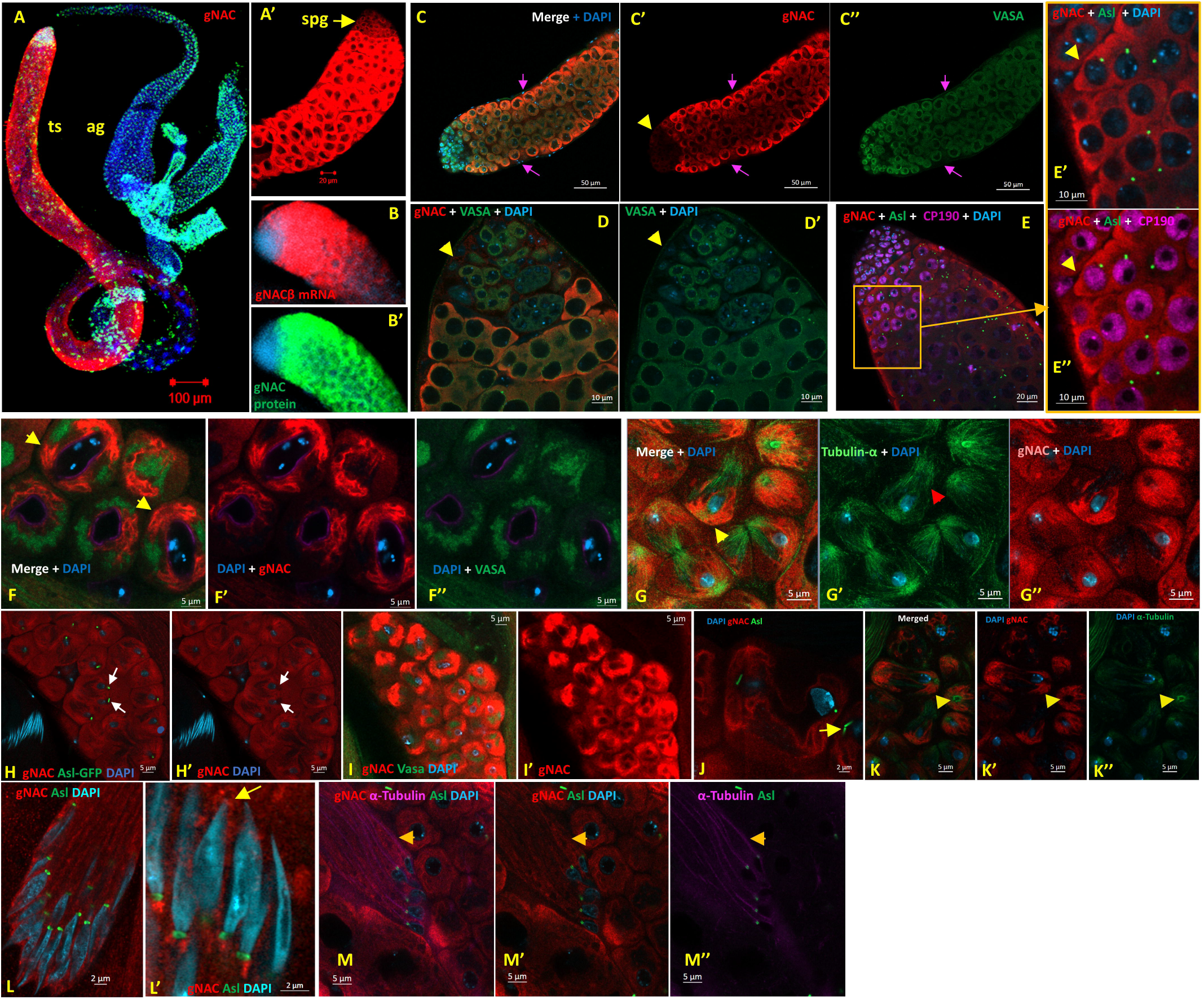
gNAC expression in testes, immunostaining by Ab against gNACβ subunit detecting the sum of gNACβ paralogs. (A) Overview of testis staining; the cytoplasm of primary spermatocytes (ts), unstained somatic accessory glands (ag), gNAC protein – red, DAPI – blue. (A’) Enlarged testis tip, brightly stained cytoplasm of spermatocytes and weakly stained spermatogonia (spg) (yellow arrow). (B, B’) FISH detection of gNACβ mRNA (red) and gNAC protein staining (green). (C-C’’) Distinct border between very weakly gNAC-stained goniablasts (yellow arrowhead) and bright staining of spermatocyte cytoplasm; unstained somatic supportive cyst cells (DAPI stained nuclei, violet arrows). (D, D’) Enlarged testis tip, stained by gNAC (red) and VASA (green). (E) Spermatocytes at the border with goniablasts, marked by Asl-GFP expression (green) and lacking gNAC staining (red) around centriole. (E’, E’’) Enlarged part of (E, orange rectangle). (F, G) Spermatocytes divisions in meiosis I, gNAC (red) and VASA (green) (F-F’’) or α-tubulin (green) (G-G’’). gNAC distribution along the centrosome microtubule asters (yellow arrowheads); separate cytoplasmic VASA associated tubulin threads compartment (red arrowhead). (H, H’) gNAC-stained round spermatids with Ast-GFP marked centrioles (green), no staining around centrioles (white arrows). (I, I’) Round spermatid with VASA (green) and gNAC (red) staining. gNAC and VASA are localized in different compartments of the cytoplasm. (J, K) Meiosis II. (J) gNAC stained cytoplasm, centriole is marked green (Asl-GFP). (K-K’’) gNAC surrounds Asl-GFP-associated centriole microtubules, seen as a circle of green spots (yellow arrowhead). (L, M) Beginning of spermatid nucleus elongation. (L), overview, centrosomes nucleating axoneme growth, gNAC is located along axoneme threads. (L’), gNAC (yellow arrow) encompasses spermatid nuclei. (M-M’’) Distal nuclear position regions of Asl-GFP marked centrosomes (green spots) known to direct the nucleation the axoneme tubulin threads; gNAC (red) is located along tubulin threads (violet, yellow arrowhead).

## 3 Discussion

Here, we present an extension of our recent results directed to the discovery of the germinal cell specific paralogs of the ubNAC known to be associated with ribosomes and has been characterized as an important partner to maintain protein homeostasis to define precise protein localization (Jomaa et al., 2022) and to control the initiation of protein translation (Zheng et al., 2022). A possibility of the specificity of the germline proteostasis is discussed while the definite general regulators of this complicated network are unknown (Dodson and Kennedy, 2020; Mercer et al., 2021). We presented the description of the developmental germline specific expression of heterodimeric gNAC, paralog of the ubiquitously expressed NAC (ubNAC), in oogenesis, spermatogenesis and early development using immunostaining to detect the sites of its locations which were followed with those of Vasa protein expression, a known marker of the germline cells. Recently we have reported that the gNAC subunits have extended IDR regions and the isoforms of its β-subunits can be differently phosphorylated (De Kegel and Ryan, 2019; Dede et al., 2020; Kogan et al., 2022).

Here we also demonstrated that CRISPR/Cas9 mediated deficiency of gNACα causes female sterility, but the ectopic expression of gNACα subunit is able to suppress the effect of a reproductive deficiency of the ubiquitous α-subunit, imitating a previously observed suppression of the lethal effect of the ubNAC β-subunit loss by the germinal β-subunit overexpression (Kogan et al., 2022). These observations prolong our considerations of crosstalk’s between the ubiquitous and germinal paralogs (Kogan et al., 2022) as a way to increase the organismal fitness by a resilience of such trait as a fertility. In accord, it has been reported recently that the ability of heterodimeric paralogs to buffer each other’s loss increased a tolerance of paralog loss (Dede et al., 2020).

Recently, we briefly mentioned the detection of gNAC in oogenesis and embryos as evidence for its presence in the oocyte germplasm and migrating embryonic germ cells (Kogan et al., 2022). Here, we traced the development of gNAC associations in the early embryonic syncytium followed by germ cell formation and their migration to the primordial somatic cells of the gonads. We unexpectedly found in the early syncytial embryos the presence of gNAC in the centrosomes that are known to be essential to organize the rapidly increasing numbers of nuclei at this stage of development (for review, (Lattao et al., 2017)). This is the case when gNAC expression is not accompanied by the presence of Vasa. The location of gNAC was also found in the centrosomes and cytoplasm of a cortex attached blastoderm cell monolayer with apically presented centrosomes situated far from the posterior embryonic pole, where the first created cells differentiate into primordial germ cells supported by inherited germplasm components. The presence of gNAC, we called as the germline specific protein, in blastoderm cells responsible for the development of somatic tissues is unexpected, but it was followed by our discovery of gNAC in the larval brain neurons (Figure 5). The observed neural association of gNAC requires for us more wider and precise researches, but brings to mind the results suggesting that many genes and gene product complexes, originally characterized for their role in the germline, also perform neural tasks (Kulkarni, 2020).

**Figure 5.**
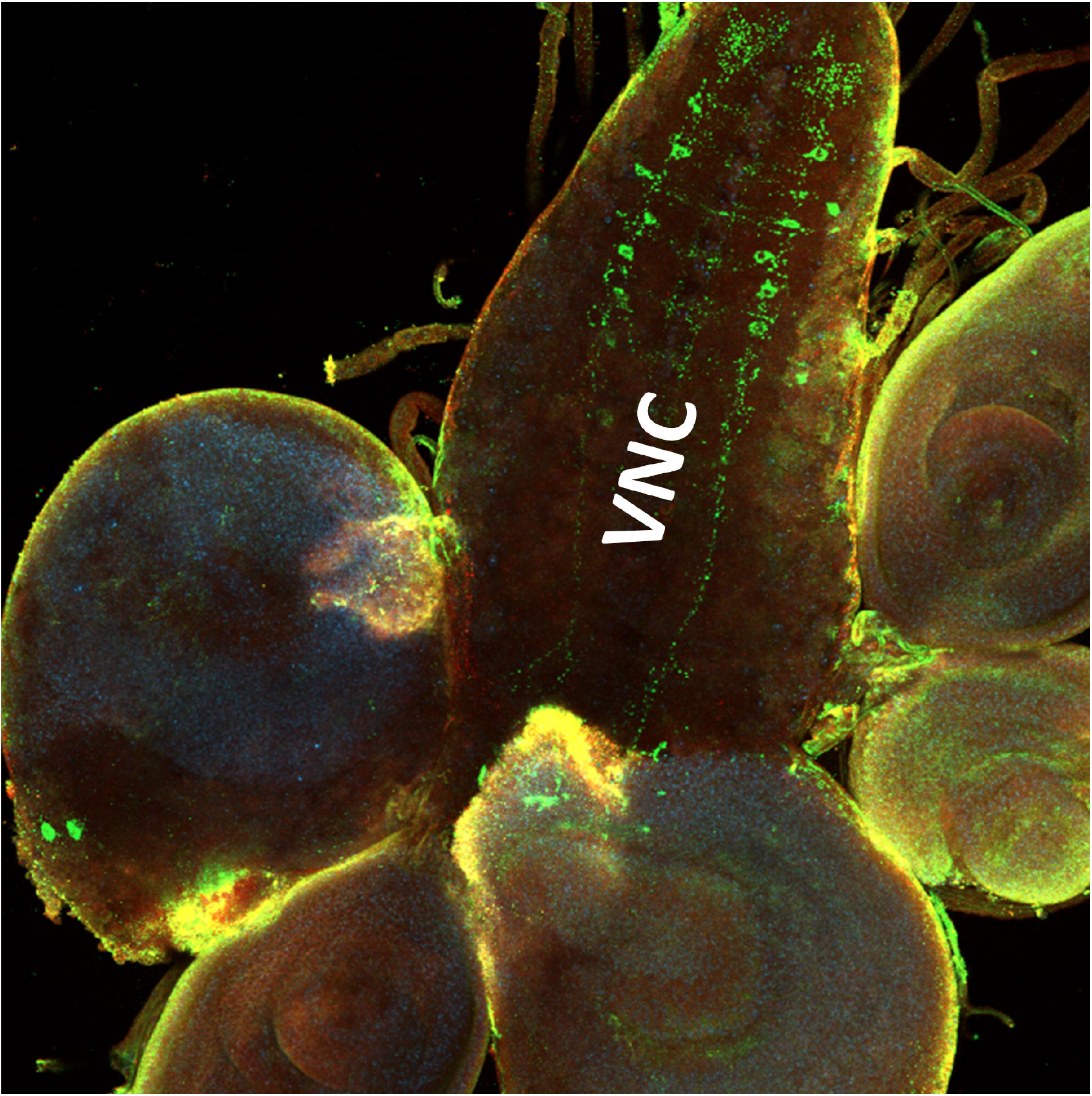
Overview of larval brain neurons (VNC, ventral nerve cord) stained for gNAC.

The presence of gNAC in the cytoplasm of pole cells has been expected, it was also preserved in the centrosomes of pole cells, the precursors of germ cells. It has been reported that the centrosome proteins participate in the organization and development of the primordial germ cells (Fang and Lerit, 2020) and centrosome nucleated microtubules facilitate germ plasm partitioning between proliferating primordial germ cells, which are highly sensitive to the dosage of germ plasm components (Lerit and Gavis, 2011).

We failed to detect accumulation of gNAC at the centrioles of growing spermatocytes but gNAC and the other PCM centrosome component, CP190, were detected around the Asl marked centriole in the male meiotic divisions where the centrosomes are known to be enlarged (Li et al., 1998). The centriole inserts into the base of the nucleus to become the basal body, which then seeds the growth of the sperm axoneme. The recent results revealing a local centrosome proteins synthesis (Bergalet and Lecuyer, 2014; Bergalet et al., 2020; Lerit, 2022) helped us to suggest gNAC involvement in centrosome proteostasis which in mammalian centrosomes is thought can be related to the functioning of mysterious, so-called centrosome satellites particles (Renaud and Bidere, 2021). At the posterior pole of the embryo, at the germ plasm location, the forming pole buds and pole cells were shown to contain cytoplasmic gNAC accumulated along microtubules of mitotic asters and as the small islands around the centriole.

We propose two functional significances of gNAC presence in the centrosomes. The first can be applied to gNAC, which can be considered as a direct component involved in the centrosome local protein synthesis, the signs of which existence have been convincingly demonstrated in a number of papers (Bergalet et al., 2020; Safieddine et al., 2021) and widely discussed (Besse and Ephrussi, 2008; Medioni et al., 2012; Chouaib et al., 2020; Lashkevich and Dmitriev, 2021; Lerit, 2022). The centrosome proteome encompassing the hundreds of proteins is proposed to use in mammals the so called centriolar satellite granules surrounding centrosome that are hypothesized as the participants of proteostasis network, but up to date their functions remain mysterious (for review, (Renaud and Bidere, 2021)). We offer to consider for *Drosophila* the direct role of gNAC paralogs in centrosome proteostasis. The second role of gNAC presence in the centrosome PCM can be related to the gNAC phoshoregulation with the help of some protein kinases populated the centrosomes. At the same time, gNAC dependent proteostasis maintaining in centrosomes can be tightly connected with various multifaceted centrosome functions including cell cycle control.

Our results regarding the selectivity of mRNA association by gNAC marked ribosomes from the oocyte hint at the existence of these ribosomes as specialized, directed to translation definite mRNAs, but this hypothesis requires in-depth study of ribosome proteins and RNA of these ribosomes. However, the topic of translation specificity realized by special ribosomes is actual (Gerst, 2018; Li and Wang, 2020; Gay et al., 2022), which is also associated with *Drosophila* ribosomes (Gershman et al., 2020; Jang et al., 2021).

In conclusion, we believe it is important to recall the possibility of the dynamic behavior in the *Drosophila* genome of gNAC isoforms equipped with elongated IDR sequences prone to phosphorylation and interaction with definite partners (Kogan et al., 2022), which may underlie its complex functions in the proteostasis and germline differentiation network. The simultaneous presence of gNAC in ribosomes and centrosomes of germ cells suggests that gNAC is a potential regulator involved in cell cycle control and modulation of carcinogenesis in the germline. Further progress in functional studies of gNAC paralogs is the discovery of its partners in the ribosome and centrosome, as well as revealing the details and functions of gNAC paralogs phosphorylation mechanisms.

## 4 Methods

### 4.1 *Drosophila* stocks

The stock *w*; endo-Asterless-GFP; MKRS/TM6B* is used for gNAC detection in the centrosomes as well as the related rapidly breeding line lacking inversions.

### 4.2 Ribosome isolation from ovaries and PCR-analysis of associated sets of mRNAs

2,000 snap-frozen *Batumi* ovaries were homogenized in a Dounce homogenizer in 1,5 ml of ice-cold lysis buffer (LB, 25mM Hepes [pH7.6], 100mM KCl, 5mM MgCl2, 1mM DTT, RNase inhibitor RiboLock (Thermo Scientific) at 40 units/ml, 0.015% digitonin, 1% NP-40, 0.5% sodium deoxycholate and 100 μg/ml cycloheximide). A protease inhibitor cocktail was used, as recommended by the company (Protease Inhibitor Cocktail Tablets, Roche). Cell fragments and mitochondria were removed by centrifugation at 12000g at 4°C for 20 min. Post-mitochondrial extract was centrifugated for 1 hour at 35,000 RPM (100000g) in a rotor Himac P50A3-0529, ultracentrifuge CP100-NX, at 4°C to get the ribosome pellet. Pellet was washed with LB buffer and dissolved in a buffer for immunoprecipitation (IPB: 50 mM Tris pH 8.0, 150 mM NaCl, 1x Complete protease inhibitor mix (Roche), 40 units/ml RNase inhibitor RiboLock (Thermo Scientific), 1% NP-40, 0.5% sodium deoxycholate). Aliquots of ribosomal extract were used for ovary total ribosome RNA preparation by the standard TRIZOL reagent (Life Technology) method and for the gNACβ immunoprecipitation (IP). Ab-gNACβ-conjugated Dynabeads Protein A (Novex by *Life* technologies) were prepared according to the manufacturers recommendation and used for immunoprecipitation. 300 μl of ribosomal extract was added to 50 μl of Ab-gNACβ-conjugated Dynabeads Protein A and incubated in a rotating vortex for 1 hour at 20 °C, then washed 5-6 times with PBS, 0,1% Tween-20. 500 μl TRIZOL reagent (Life Technology) was added to magnetic particles and thoroughly mixed, 1/10 volume CHCl_3_ was added, left for 10 minutes on ice and centrifuged at 10,000 RPM for 5 minutes at 4°C. 1 μg of glycogen was added to the aqueous phase and RNA precipitated by adding an equal volume of isopropanol and incubating overnight at −20 °C, washed with 70% ethanol, dissolved in DEPC treated water and quantified with a NanoDrop 1000 spectrophotometer (Thermo Scientific).

### 4.3 Quantitative real-time PCR (q-RT-PCR)

q-RT-PCR primers were designed using Primer 3. cDNA for qPCR was synthesized using random primers and the First Strand cDNA synthesis kit (MINT Reverse Transcriptase, Evrogen, Russia) according to manufacturer’s instructions. The relative abundance of transcripts was estimated using quantitative real-time PCR with Hot Start DNA polymerase (Evrogen, Russia) and SYTO@ 13 green fluorescent nucleic acid stain (Life Technologies) on the DT-96 instrument (DNA Technology, Russia). The results were normalized to the amount of the *Rpl32* mRNA. The primers used in q-RT-PCR quantifications are listed in the Supplementary Table S1.

### 4.4 Constructs of plasmids and CRISPR mutagenesis

To delete the *CG4415* gene from the stock BDSC #55821 (*y*^*1*^ *M{GFP[E*.*3xP3]=vas-Cas9*.*RFP}ZH-2A w*^*1118*^), the unique guide RNAs directing Cas9 to the regions upstream and downstream of the gene were selected with Target Finder (Gratz et al., 2014). Synthesized oligonucleotides for the guide RNAs (guide_*CG4415*_up_s /guide_*CG4415*_up_as and guide_*CG4415*_down_s/guide_*CG4415*_down_as, Table S2) were annealed and cloned into the *gRNA* plasmid *pU6-BbsI-chiRNA* (addgene66 #45946) linearized with *Bbs*I. Homology-directed repair (HDR) donor plasmid was generated on the basis of the *pDsRed-attP* backbone (addgene#51019). Left homology arm (LA), corresponding to the 1 kb region upstream of the *CG4415*, was PCR amplified with the oligonucleotides *up_as_XhoI* and *up_s_EcoRV* (Table S2) using genome DNA of the BDSC #55821 line as a template. PCR-fragment was digested with *Xho*I and *Eco*RV restriction endonucleases and cloned into the corresponding sites of the *pDsRed-attP* plasmid. Right homology arm (RA) corresponding to the 1 kb region downstream *CG4415*, was PCR amplified with the *down_s_NheI* and *down_as_SmaI* primers using the same template. PCR fragment was digested with *Nhe*I and *Sma*I restriction endonucleases and cloned into the corresponding sites of *pDsRed-attP*, containing LA. The mix of the two guide RNAs and the donor plasmid was injected into the BDSC #55821 embryos. The germ-line transformation of the embryos was performed according to the standard protocol (Rubin and Spradling, 1982). Founder males were individually crossed with *yw*^*67c23*^, +*/CyRoi* females to generate the progeny with the *CyRoi* balancer chromosome, lacking the *vas-Cas9* -GFP marker and the *CG4415* gene substituted by *dsRed*-containing fragment from the donor vector. Two lines with moderate-to-increased high levels of *DsRed* expression in eyes were created and designated hereafter as *mDsRed* and *incDsRed*, respectively. DNA from transgenic flies was extracted according to standard proteinase K treatment – phenol-chloroform extraction – ethanol precipitation protocol. Deletions were checked by PCR with one primer to deleted *CG4415* gene and the another one to the flanking region.

The organization of the genome insertions *mDsRed* and *incDsRed* was also checked by MinIon sequencing. Libraries were prepared from 400 ng of genomic DNA according to the protocol for Rapid Sequencing Kit (SQK-RAD004, ONT). Libraries were loaded into a MinIon R 9.4.1. flowcell and sequenced without basecalling. Approximately 1 Gb of data were obtained in each case. A set of .fast5 files from the sequencing runs was basecalled on a standalone GPU-enabled server with Guppy Version 3.5.2 using the dna_r9.4.1_450bps_hac profile. Reads with Q<7 were filtered out in the course of basecalling. The resulting FASTQ files were loaded to the local Galaxy instance for quality checks and analysis. Nanostat (https://github.com/wdecoster/nanostat) tool in the Galaxy showed that ∼1 Gb of reads with N50= 10780 was obtained. Adapters were trimmed by Porechop (https://github.com/rrwick/Porechop) using default settings (reads with a middle adapter were split). Processed reads were mapped to the r6.22 release of the *D. melanogaster* genome using Minimap2 software with the Oxford Nanopore read-to-reference mapping profile (minimap2 -x map-ont).

For the both lines, a gap in a coverage in the region of planned replacement (*CG4415*, chr2L:1,036,879-1,038,774, Figure S1) was found. To check the structure of the insertions, processed reads were converted to FASTA using the FASTQ-to-FASTA converter in Galaxy (Blankenberg et al., 2010) and BLASTed (dc-megablast with the default settings) against nr BLAST database (https://blast.ncbi.nlm.nih.gov/Blast.cgi). BLAST output was converted to BED file for visualization in IGV. It was found that the CRISPR recombination in the case of *incDsRed* produces rearranged genome region consisting of two full and one partial repeat of the main part of the plasmid used for injection (Figure S1), including 3 copies of the *dsRed* gene. *mDsRed* genome insertion arranged as expected with single copy, in proper orientation and position.

### 4.5 Embryo and tissue staining and confocal imaging

0-1-h embryos were collected, dechorionated, devitellinized and immunostained as previously described (Kogan et al., 2022). Testes and ovaries of adult (1-3 day old) males and (5-7 day old) females were dissected in phosphate-buffered saline (PBS) at 4°C, washed with PBT, fixed in 3,7% formaldehyde in PBT for 30 min at RT and then processed and immunostained, like embryos. The following primary antibodies were used: rabbit polyclonal anti-gNACβ (1:500, developed in our laboratory (Kogan et al., 2022), rat anti-VASA 1:200 (DSGB: AB_760351), mouse anti-α-tubulin 1:200 (T9026, Sigma-Aldrich). IgG Alexa Fluor 633-conjugated goat anti-rabbit (1:500, Invitrogen, USA), Alexa Fluor 546–conjugated goat anti-rat (1:500, Invitrogen, USA) and anti-mouse IgG Alexa Fluor 488 (1:500, Invitrogen, USA) secondary antibodies were used. Confocal microscopy was performed on ZEISS LSM 900 system (Zeiss, Germany).

## Author Contributions

Elena Mikhaleva is responsible for confocal imaging, Natalia Akulenko performed CRISPR-CAS mutagenesis and plasmid construction, Oxana Olenkina created transgenic flies, Yuri Abramov is responsible for fly genetics and crosses, Sergey Lavrov performed genomic sequencing and analysis, Toomas Leinsoo and Galina Kogan obtained qPCR quantifications data, Vladimir Gvozdev is a conception author. Vladimir Gvozdev and Galina Kogan are performed paper arrangement.

## Acknowledgements

We are grateful to Professors D.M. Glover (Caltech, US) and J.W. Raff (Oxford, UK) for their kind gifts of *Drosophila* stocks expressing *Asl-GFP* constructs

## Data Availability Statement

The original contributions presented in the study are included in the article/supplementary material, further inquiries can be directed to the corresponding author/s.

