## Supplementary materials for "A study of the expression of germline-specific paralogs of the ubiquitously expressed NAC complex revealed their association with centrosomes"

### 1 Supplementary Figures and Tables

#### 1.1 Table S1. Primers for qRT- PCR

| **FlyBaseID** | **Gene** | **Direction** | **Sequence (5’→3’)** | **PCR product, b.p.** |
| --- | --- | --- | --- | --- |
| FBgn0000042 | **Act** | forward | GCAGTGCACCGGAAAGAAAG | 189 |
|  | CG4027 | reverse | CATCCGAGCGCCAAATCAAG |  |
| FBgn0000055 | ***Adh*** | forward | AAACTGGCCCCCATTACCG | 216 |
|  | CG3481 | reverse | CAAGTCCAGTTTCCAGATG |  |
| FBgn0262167 FBgn0262167  FBgn0262167 FBgn0262167 FBgn0262167 | ***ana1*** | forward | CCACGTAACCAGCCATCAGT | 186 |
|  | CG6631 CG6631CG CG6631 | reverse | AGCCTGTTCTGGAATCGGTG |  |
| FBgn0027513 FBgn0262167  FBgn0262167 FBgn0262167 FBgn0262167 | ***Ana2*** | forward | TCCAACGCAGATGTCCTCAC | 149 |
|  | CG8262 CG6631CG CG6631 | reverse | GATACCAGAGCCGCCAGATC |  |
| FBgn0266111 | ***ana3*** | forward | TTGTCCTGGGTATCGTTGGC | 139 |
|  | CG13162 | reverse | GCATCATTTCGTGGGCATCC |  |
| FBgn0261004 | ***asl*** | forward | CCCATCACATCCGCACCATA | 190 |
|  | CG2919 | reverse | TAGCTCAGCCTGCATGATGG |  |
| FBgn0003884 | ***αTub84B*** | forward | CAACTCCATCCTGACCACCC | 137 |
|  | CG1913 | reverse | TCAGACGGTTCAGGTTGGTG |  |
| FBgn0000182 | ***bicC*** | forward | GGACGACACGCATACCCATA | 151 |
|  | CG4824 | reverse | GAAGGAGATGAGCAGCGGTG |  |
| FBgn0013765 | ***cnn*** | forward | AACCCTGCAATCCCAGCTAC | 159 |
|  | CG4832 | reverse | TCCAGGGATCGTTGCAGTTC |  |
| FBgn0000283 FBgn0000283 FBgn0000283 | ***Cp190*** | forward | TAAGCAGCAATCCCCACAGG | 187 |
|  | CG6384 | reverse | GTGGCCCTTTACAATGTGCG |  |
| FBgn0000405 | ***CycB*** | forward | ACAGAAGGAGGTGTCCCACAAGATG | 204 |
|  | CG3510 | reverse | TCCTCGTACTTGGTGGCTATGAAGAG |  |
| FBgn0260634 | ***eIF4G2*** | forward | CCATCGGATTGCAGAGAAACAAATA | 356 |
|  | CG10192 | reverse | CGTGTTGTGATACAAATGAAATGGA |  |
| FBgn0023515 | ***Gcna*** | forward | GGGCAAATACGGAGGACACA | 115 |
|  | CG14814 | reverse | GACAGGAGCTCTACATCGCC |  |
| FBgn0030566 | ***gNACβ*** | forward | AATCGTGTTCGTCCTCGCAA | 166 |
|  | CG18313 | reverse | CCGATGGCACCTTCTCATTATCCG |  |
| FBgn0031296 | ***gNACα*** | forward | GCCCAATGTGTTCAGTCTGC | 165 |
|  | CG4415 | reverse | GTACCGAGTTGACCCTGCTC |  |
| FBgn0005695 | ***gcl*** | forward | ACGGTGTGGTCGGAGTCAAG | 213 |
|  | CG8411 | reverse | CGTAGTCGGGATGCAGTCGT |  |
| FBgn0001120 | ***gnu*** | forward | TCACTCCCCTCTCCACCGAA | 188 |
|  | CG5272 | reverse | AAATCGACGGGGCAAAGTGG |  |
| FBgn0004419 | ***me31B*** | forward | ACAAGTGACGATATGGGCTGG | 190 |
|  | CG4916 | reverse | CTCGAATTCATTGCCTCGCG |  |
| FBgn0000228 | ***Nin*** | forward | AGAACACGGAGCTTGAGTCG | 174 |
|  | CG14025 | reverse | TCTCCTCCTCCTTGTCCTCG |  |
| FBgn0002962 | ***nos*** | forward | ACGCTTCGCAGTTGTTTCAA | 190 |
|  | CG5637 | reverse | ATCGCGCACTCTACTTTCCA |  |
| FBgn0003015 | ***osk*** | forward | CCAGCAAGCCAGCGTAAATG | 125 |
|  | CG10901 | reverse | GGCTTTGGGTTCTGCAGCTT |  |
| FBgn0016053 | ***pgc*** | forward | TGGCATCCTACGACAATGGA | 80 |
|  | CG32885 | reverse | CTCATTCATCTCCCGCTCCC |  |
| FBgn0004872 | ***piwi*** | forward | CCGTGGGGTGACCAATATGAT | 181 |
|  | CG6122 | reverse | GGACACGAGGATTCTCGATGC |  |
| FBgn0086690 | ***Plp*** | forward | ACGAGAACTTCACTGGCGAG | 180 |
|  | CG33957 | reverse | CCAGTCGTTTCCTCATGGCT |  |
| FBgn0000826 | ***png*** | forward | GAGAGGAAGGTGTGCGTGAA | 173 |
|  | CG11420 | reverse | TTGGGCACGTACTCCATCAC |  |
| FBgn0024733 | ***RpL10*** | forward | AGTGAAGCTTTGGAAGCTGGACGCA | 137 |
|  | CG17521 CG17521 CG17521 | reverse | CCAGCGCACGACAACATTTTGTTG |  |
| FBgn0002626 | ***RpL32*** | forward | ATGACCATCCGCCCAGCATAC | 88 |
|  | CG7939 | reverse | GCTTAGCATATCGATCCGACTGG |  |
| FBgn0039731 | ***Sas-6*** | forward | GCTGCCGAGGTCCATCAATA | 127 |
|  | CG15524 | reverse | TCTCCAACGAAGCCTTGTCC |  |
| FBgn0003514 | ***sqh*** | forward | ACCTCCAATGTGTTCGCCAT | 109 |
|  | CG3595 | reverse | GCAGATCCTCCTTCTCGACG |  |
| FBgn0003891 | ***tud*** | forward | CGTGGGTCCGTATCTGAAGG | 213 |
|  | CG9450 | reverse | CGTATCCGCCTGTACTCCAC |  |
| FBgn0283442 | ***vas*** | forward | TTGGAAGACCCCAGGTAGTG | 208 |
|  | CG46283 | reverse | CGATCCACGAAATCCAGAAG |  |
| FBgn0041087 | ***wun2*** | forward | TTTAGGGGACAGGACAAGCG | 196 |
|  | CG8805 | reverse | GAGGTCTCAGCCGTCCAATG |  |

#### 1.2 Table S2. Primers used for constructs of plasmids and CRISPR mutagenesis

| **Oligonucleotide name** | **Oligonucleotide sequence** |
| --- | --- |
| *guide_CG4415_up_s* | CTTCGAACTGTGCACTTAATTTCG |
| *guide_CG4415_up_as* | AAACCGAAATTAAGTGCACAGTTC |
| *guide_CG4415_down_s_* | CTTCGTACAAAGGGGTTCTTCAGT |
| *guide_CG4415_down_as* | AAACACTGAAGAACCCCTTTGTAC |
| *CG4415_up_as_XhoI* | TTATAACTCGAGTGCTGCAGCGCCTCGCAAATT |
| *CG4415_up_s_EcoRV* | TCTCCGGATATCCGAGTTCCAAGTCCGCGCAGAA |
| *CG4415_down_s_NheI* | CCCCTTGCTAGCGCGAAATTGCCATCTTATTTCA |
| *CG4415_down_as_SmaI* | TCGTTCCCCGGGACAAGTAGTTCATTTCGAGAAA |

#### 1.3 Supplementary Figure S1

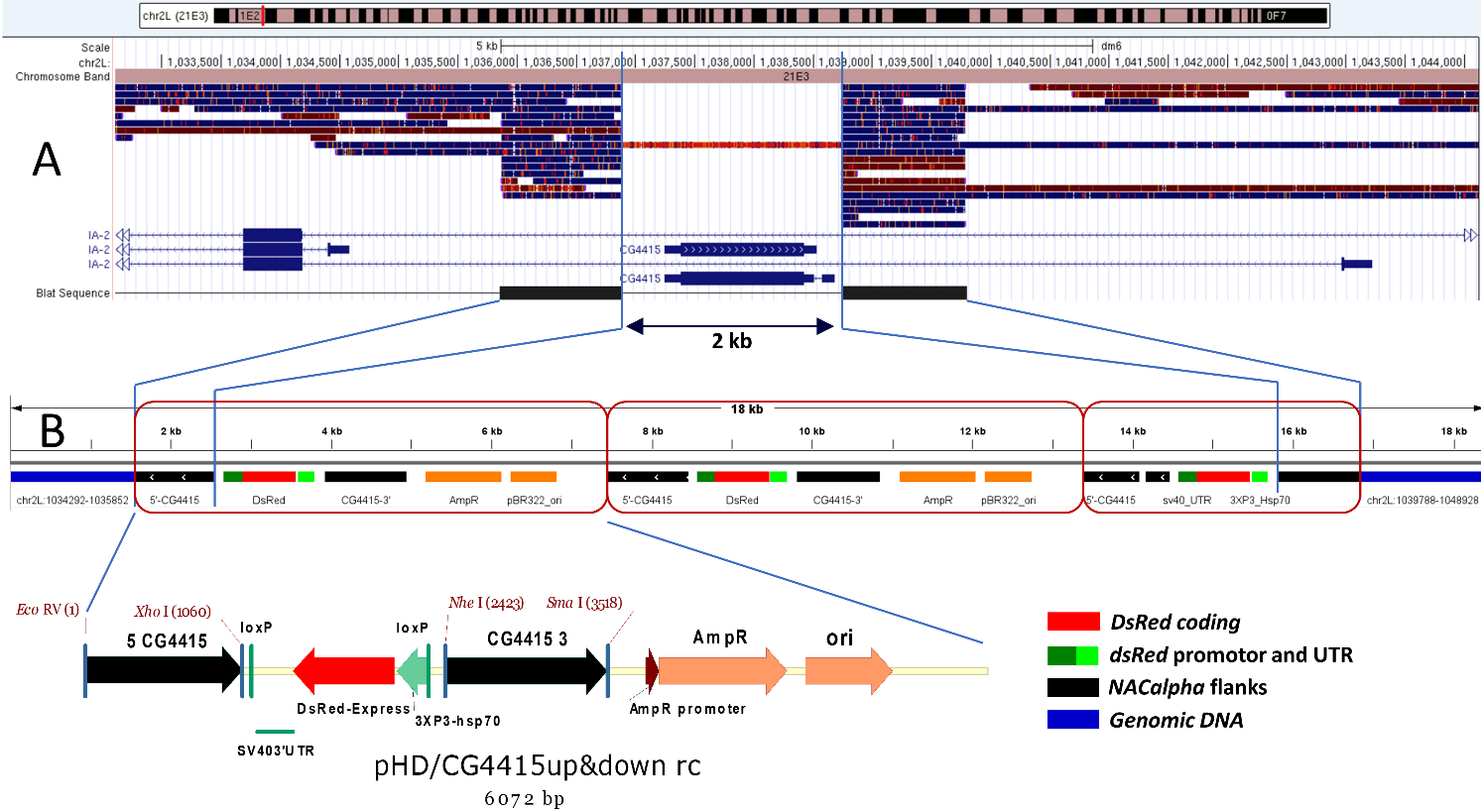

**Figure S1.** Structure of the *incDsRed* insertion replacing *CG4415* gene

**(A)** Alignment of Nanopore reads from *insDsRed* transgenic line to the *chr2L:1,033,106-1,044,320* genome region encompassing the *CG4415* gene*.* **(B)** The structure of the *insDsRed* insertion. *CG4415* gene region of 2 kb in size was replaced by the two full and one partial copy of the transformation vector pHD/CG4415up&down (marked by brown ovals). The genomic regions used for the replacement induction (5’-CG4415 and CG4415-3’) are shown in black, the other components of the vector stained red (*DsRed*), green (regulatory regions of *DsRed*) and orange (plasmid parts).
